# Comprehensive Codon Usage Analysis Across Diverse Plant Lineages

**DOI:** 10.1101/2023.11.20.567812

**Authors:** Aasim Majeed, Wahid Ul Rehman, Amitozdeep Kaur, Sreemoyee Das, Josepheena Joseph, Amandeep Singh, Pankaj Bhardwaj

## Abstract

The variation of codon usage patterns in response to the evolution of organisms is an intriguing question to answer. The purpose of this study was to investigate the relevance of the evolutionary events of vascularization and seed production with the codon usage patterns in different plant lineages. We found that the optimal codons of non-vascular lineages generally end with GC, whereas those of the vascular lineages end with AU. Correspondence analysis and model-based clustering showed that the evolution of the codon usage pattern follows the evolutionary event of the vascularization more precisely than that of the seed production. The dinucleotides CpG and TpA were under-represented in all the lineages, whereas the dinucleotide TpG was found over-represented in all the lineages, except algae. Evolutionary-related lineages showed similar codon pair bias (CPB). The dinucleotide CpA showed a similar representation as those of its parent codon pairs. Although natural selection predominates over mutational pressure in determining the codon usage bias (CUB), the relative influence of mutational pressure is higher in the non-vascular lineages than in the vascular lineages.

## Introduction

The variation of codon usage patterns in response to the evolutionary events of plants, such as vascularization and seed production, is an intriguing question to answer. With respect to vascularization, the plants can broadly be categorized into non-vascular (algae and bryophytes) and vascular (pteridophytes, gymnosperms, and angiosperms) lineages. With respect to seed production, the plants can be categorized into seedless (algae, bryophytes, and pteridophytes) and seeded (gymnosperms and angiosperms) lineages. The modern-day seed plants are represented by gymnosperms comprising four different lineages (cycads, conifers gnetophytes, and *Ginkgo biloba*) and angiosperms comprising dicots and monocots. Fossil records showed considerably greater seed plant diversity, dating back to the Late Devonian approximately 360 million years ago (Wang et al. 2018). Vascular plants were thought to have evolved ∼410 million years ago. They diverged into several lineages, only two of which survive today: the euphyllophytes (ferns and seed plants) and the lycophytes (club mosses and spike mosses) (Banks et al. 2011). Algae represent a paraphyletic diverse group of photosynthetic eukaryotes. The algal groups, chlorophyta and streptophyta together with land plants (embryophyta) constitute viridiplantae (Ruhfel et al. 2014).

The occurrence of 64 triplet codons coding for 20 amino acids implies that certain synonymous codons are used more frequently in the genome/genes, a phenomenon termed as codon usage bias (CUB). The major factors which govern the CUB include mutational pressure and natural selection, the equilibrium of which is important for codon usage in both prokaryotes and eukaryotes (Bulmer 1991; Hershberg and Petrov 2008). Some mutations may occur more frequently than expected and this mutational bias may lead to changes in allele frequency, a phenomenon known as mutational pressure. Neutral mutations at degenerate codon positions create diverse synonymous codons coding for same amino acid. These neutral mutations spread and get fixed in a population independent of selection. If these neutral mutations are biased, the resulting mutational pressure would ensure enhanced frequency or possible fixation of some codons over others. In absence of selection, this mutational pressure can determine codon usage (Peden 1999). A high level of any of the four nucleotides (Zhong et al. 2007) or a high GC-content at the third position of codons (Sueoka 1988) suggests involvement of mutational pressure governing the CUB. In general, the CUB in mammals and prokaryotes having high GC- or AT-contents is driven by mutation, whereas, in plants and Drosophila, the translational selection is the driving force (Uddin 2017). Translation of superior codons occurs under the balance of mutation, drift, and selection (Behura et al. 2010). Highly expressed genes exhibit a strong bias in which selection turns against rarely used codons and in the favor of frequently used ones. Although different approaches exist for the determination of CUB pattern, the Relative Synonymous Codon Usage (RSCU) has gained wide popularity. Besides the natural selection and mutation pressure, the CUB is determined by several other factors. The expression level, the position of codons, gene length, GC-content, hydropathicity, RNA stability, and tissue type may also impart their influence on the pattern of codon usage (Powell and Moriyama 1997; Akashi 1997; Carlini and Stephan 2003; Powell et al. 2003; Plotkin et al. 2004). The patterns of codon usage provide useful insights into the prediction, classification, and evolution of genes, host-pathogen co-evolution, and adaptation (Wang et al. 2018). The phylogenetic distance may also contribute to the divergence of CUB (Zhao et al. 2016).

CUB is widespread across genomes and may have a profound role in genome evolution. In genome biology, the genome-wide codon usage pattern, its causes and consequences, and the identification of selective forces shaping the evolution of the codon usage patterns are the important realms of study (Behura and Severson 2013). Different studies have analyzed the codon usage pattern in isolated or a few plant species such as *G. biloba* (He et al. 2016), *T. contorta* (Majeed et al. 2020), several pteridophytes species (Das et al. 2019), *Oryza sativa*, *Zea mays*, *Triticum aestivum*, *Hordeum vulgare*, and *Arabidopsis thaliana* (Kawabe and Miyashita 2003; Liu and Xue 2005a), *Cephalotaxus oliveri*, *Gnetum parvifolium* (Qi et al. 2015). Studies evaluating and comparing the genome-wide codon usage pattern of different plant lineages are lacking. In the evolutionary history of plants, vascularization and seed production are important evolutionary events. The divergence of codon usage varies among the species as well as genes (Dohra et al. 2015). Apart from the species and genes, it is quite intriguing to evaluate how the codon usage pattern varies among different plant lineages. Moreover, it is fairly enthralling to assess the relative influence of vascularization and seed production on codon usage pattern. This study was therefore undertaken to investigate the variations in the general pattern of codon usage; to evaluate the relative impact of the factors influencing the CUB in different plant lineages and to evaluate the relation of vascularization and seed production on codon usage.

## Material and methods

### De novo assemblies and identification of open reading frames (ORFs)

The paired-end raw reads of 68 species belonging to algae, bryophytes, pteridophytes, gymnosperms, dicots, and monocots were retrieved from the NCBI SRA database (Table 1). The *de novo* assemblies of the individual species were generated through Trinity v1.6 (Grabherr et al. 2011). CD-HIT-EST v4.6 (Li and Godzik 2006) was used to remove the sequence redundancy using the sequence identity cuttoff value of ≥ 0.9. The full-length sequences of the resulting longest transcripts were identified through TRAPID (Van Bel et al. 2013) using Plaza database as a reference. The ORFs of these full-length transcripts were then identified by Transdecoder using default settings. These coding sequences (CDS), containing a start and a stop codon, were used for the analysis of codon usage patterns.

**Table 1:**
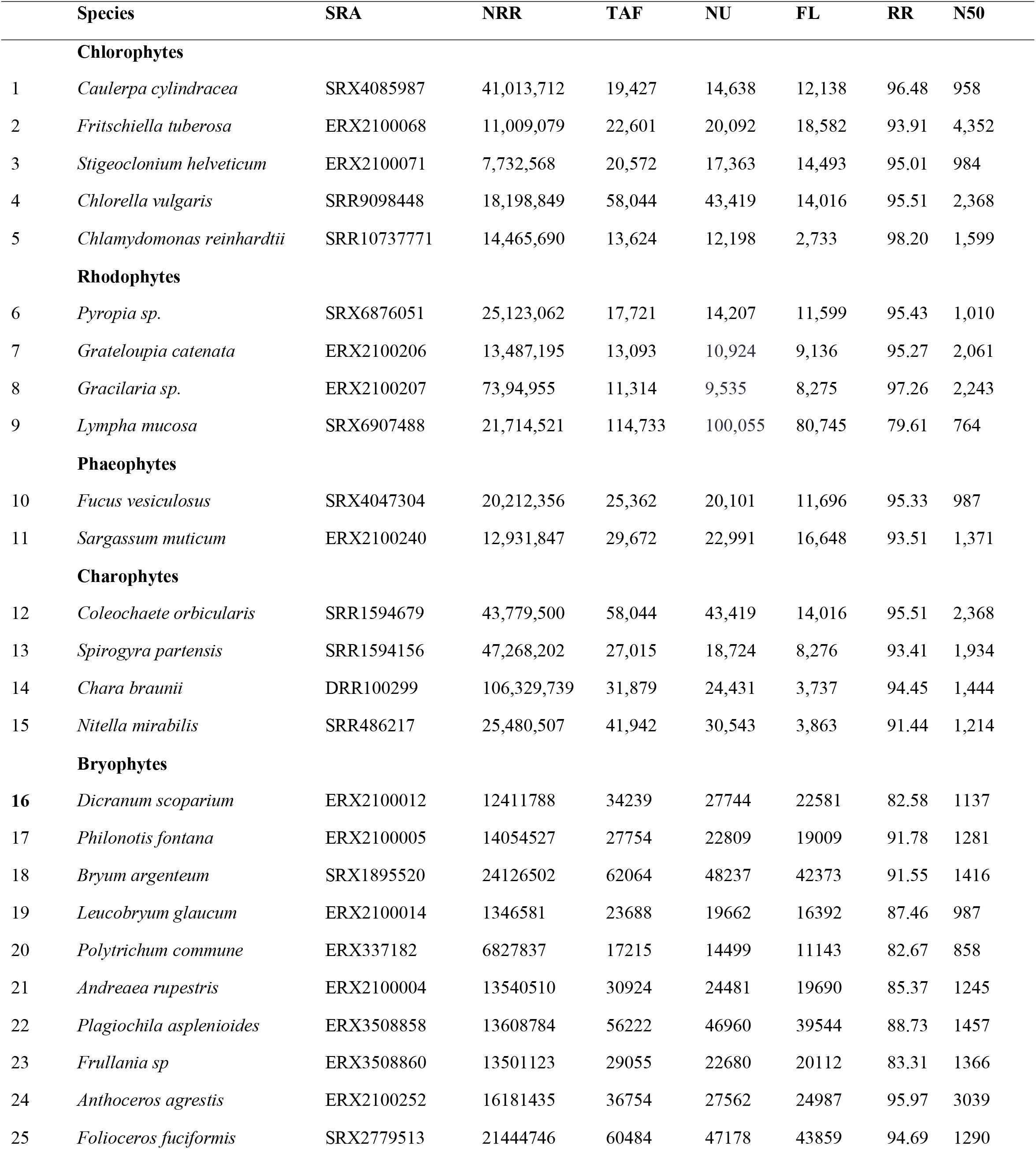
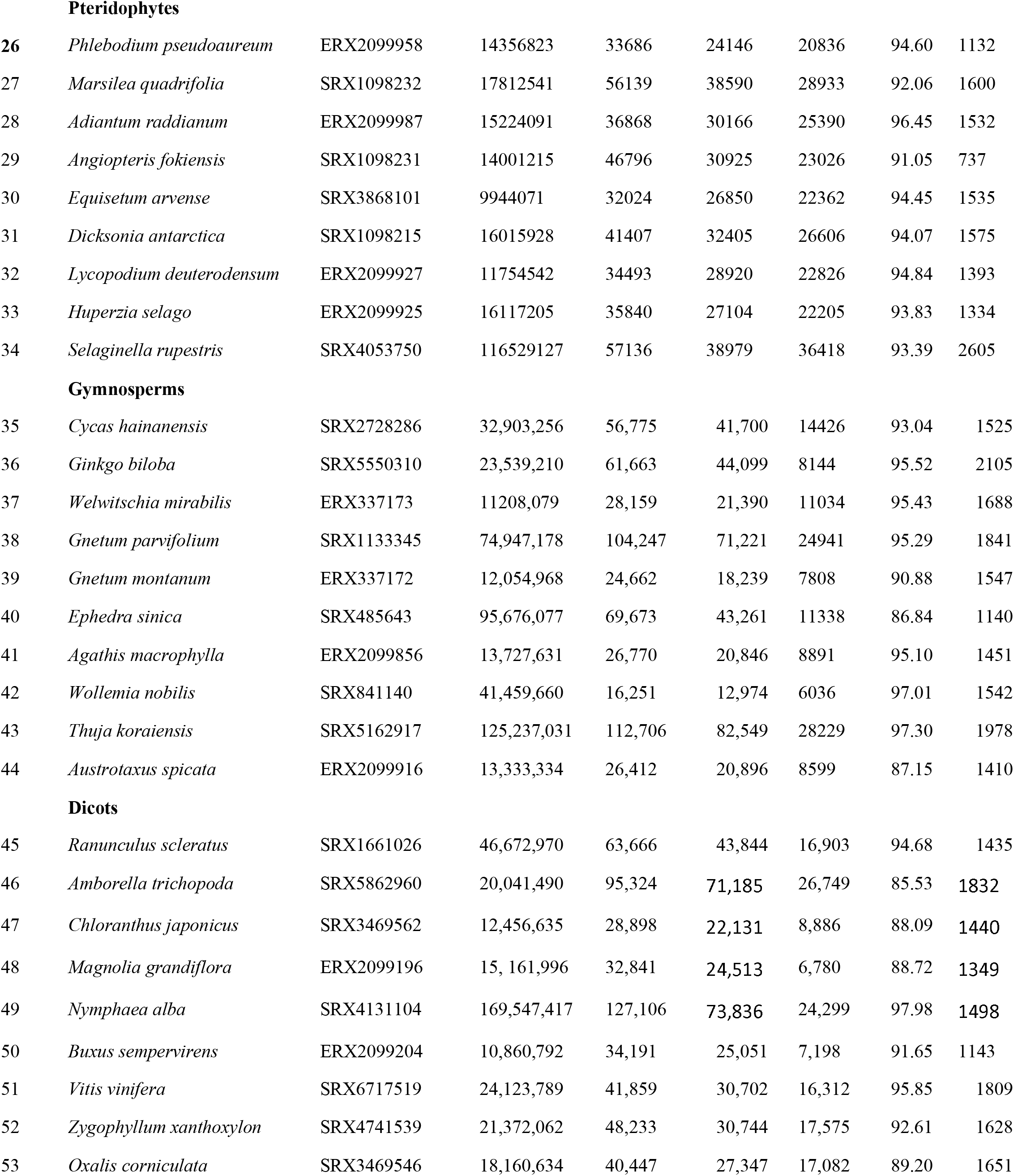
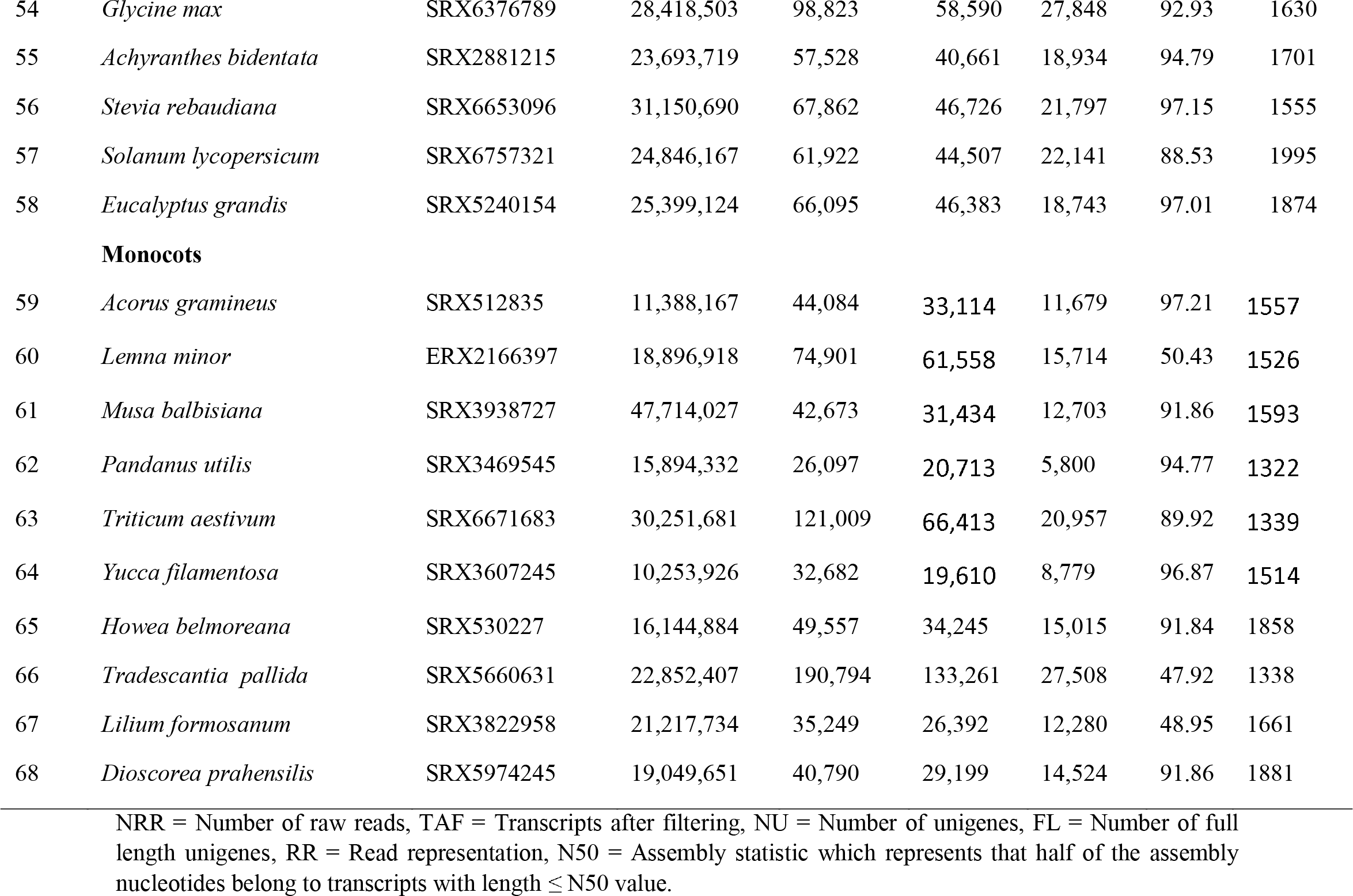
Details of individual assemblies.

### Determination of the codon usage indices and compositional properties

CodonW v1.4.2 (Peden, 2005) was used for the determination of overall nucleotide composition and composition at the third position of the codons using default settings. The codon bias indices like Relative Synonymous Codon Usage (RSCU) and Effective Number of Codons (ENC) were determined as described in Majeed *et al*., (2020). ENC is an index of CUB whose value ranges from 20 (perfect bias) to 61 (equal usage of degenerate codons). For the evaluation of GC-content at 1^st^, 2^nd^, and 3^rd^ codon positions, we employed a Perl script (https://github.com/hxiang1019/calc_GC_content.git).

### Determination of the optimal and the avoided codons

An optimal codon is a codon that is significantly more frequently used in putatively highly expressed genes, than those with lower levels of expression. It must be noted that a frequent codon may not necessarily be an optimal codon (Peden, 1999). We determined optimal codons using CodonW, which uses the upper and lower 5% genes on the principle axis, generated through correspondence analysis, as the two contrasting datasets of high and low expression, respectively. The expression estimates are predicted indirectly by codonW through codon usage indices. Then the CodonW estimates significance using a two-way chi-squared contingency test on the codon pairs from these two groups of genes, with a p-value cutoff of <0.01 (Peden, 1999). The RSCU is a measure of codon usage bias. RSCU value of 1 denotes no bias. The codons having RSCU >1 are biased and are therefore preferred over other codons, regardless of the gene expression. The codons having RSCU value <1 are used less frequently (Goldman and Yang 1994; Guan et al. 2019). Further, RSCU < 0.75 represent the avoided codons (Guan et al. 2018). We determined the avoided codons as codons having RSCU <0.75.

### Correspondence (CA) and Clustering Analysis

The CA is used to explore relationships among qualitative/categorical variables. Like principal component analysis, it provides a solution for summarizing and visualizing data set in two-dimension plots. To summarize and visualize the synonymous codon usage variation among different plant lineages, we performed CA on their RSCU values using the FactoMineR and factoextra packages in R. A contingency table consisting of 59 columns, each corresponding to the RSCU value of one sense codon, and rows corresponding to genes was used as input. To gain more insight into the pattern of variation of codon usage in different plant lineages, we used a model-based clustering analysis of RSCU values of the codons, using an R package, mclust.

### Determination of the relative impact of mutation and selection on CUB

We used a neutrality plot analysis to determine the relative influence of the mutation and the selection on the CUB. In the neutrality plot analysis, the relation between the GCs at the position 1 and 2 (GC12) and GC at the position 3 (GC3) of the codons were assessed. If GC12 and GC3 are statistically correlated and the regression slope is close to one, the mutational pressure is the driving force. This is because mutational bias affects all codon positions equally, resulting in a strong correlation between GC12 and GC3, thereby increasing the slope of regression line. As the slope approaches more horizontal, the selection turns to be a dominant force (Wu et al. 2015). Selection is well-known to drive non-synonymous substitutions, it also acts more strongly on the third codon position, which is more variable and less constrained than the first two positions (Du et al, 2014). In both cases, it results in a weaker correlation between GC12 and GC3, hence more horizontal regression line. Selection on synonymous codons, arising due to variations at 3^rd^ codon position seems to be influenced by post-transcriptional and translational pressures to ensure translational accuracy, proper protein folding, and/or mRNA stability (Du et al, 2014).

### Evaluation of dinucleotide bias

Codon usage bias may be influenced by dinucleotide bias. We estimated the mononucleotide and the dinucleotide frequencies using Simmonic sequence editor, version 1.7 (Simmonds 2012). Then the dinucleotide bias was evaluated as the ratio between the observed frequency and the expected frequency (odd ratio) of the 16 possible dinucleotides. The expected frequencies were calculated by multiplying the frequencies of the two nucleotides of the dinucleotide (Kapoor et al. 2010). Under no dinucleotide bias, the odd ratio i.e. pxy = fxy/fxfy shall be equal to 1. Any deviation from 1 depicts bias, with pxy > 1.23 indicating overrepresentation and pxy < 0.78 indicating underrepresentation (Karlin and Mrázek 1996).

### Evaluation of Codon Pair Bias (CPB)

Independent of codon bias, juxtaposed codons in open reading frames (ORFs) appear to be non-random, with some codon pairs used more or less frequently than expected from their frequencies (Coleman et al. 2008; Kunec and Osterrieder 2016), a phenomenon termed as codon pair bias (CPB). The CPB may also affect the codon usage pattern of a species. The determination of the CPB was performed by calculating the Codon Pair Scores (CPSs) according to Coleman *et al.,* (2008). CPS is the the natural logarithm of the ratio of the observed to the expected number of a specific codon pair in all the sequences and is calculated as:

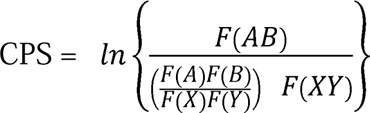

Where AB is the codon pair coding the amino acid pair XY and F is the counts of each factor (A, B, AB, X, Y, and XY). A negative value of CPS indicates that the codon pair is underrepresented, whereas a positive value reveals that the codon pair is overrepresented (Kunec and Osterrieder, 2016). CPS was calculated for each of the 3721 possible codon pairs (61 × 61 codons) using the R package, CPBias. Codon Pair Bias (CPB) scores were then calculated as the mean of the CPS values of all codon pairs.

## Results

### Identification of ORFs

A total of 1643.912 million paired-end raw reads, belonging to 68 species, were processed and *de novo* assembled into 6,873,001 transcripts, constituting 68 individual assemblies. Filtering of the transcripts <500 bp lengths resulted into 3,077,232 sequences. These sequences were subjected to removal of sequence redundancy, which generated 2,229,137 transcripts. A total of 1,197,256 transcripts were identified as full length sequences, which contain a start and a stop codon. The ORFs of these full-length sequences were then used for the analysis of CUB. The detailed information about the individual assemblies of 68 species is presented in Table 1.

### GC compositional properties

The mean GC-content in algae, bryophytes, pteridophytes, gymnosperms, dicots, and monocots was observed to be 0.55, 0.51, 0.47, 0.45, 0.38, and 0.49 respectively. The difference of mean GC-content and mean GC3-content between seedless (mean GC = 0.521, mean GC3 = 0.546) and seeded (mean GC = 0.432, mean GC3 = 0.422) lineages was found to be 0.089 and 0.124 respectively. Similarly, for vascular and non-vascular lineages, the respective mean GC-content of 0.440 and 0.539 with a difference of 0.099 and the respective mean GC3-content of 0.429 and 0.580 with a difference of 0.151 was observed. Thus, the variation of GC-content, as well as GC3-content, appeared slightly more pronounced for the divergence of plants with respect to the evolutionary event of vascularization than that of seed production. This was statistically corroborated by Kruskal-Wallis test followed by Dunn test as post hoc analysis (Figure 1A, 1B).

**Figure 1:**
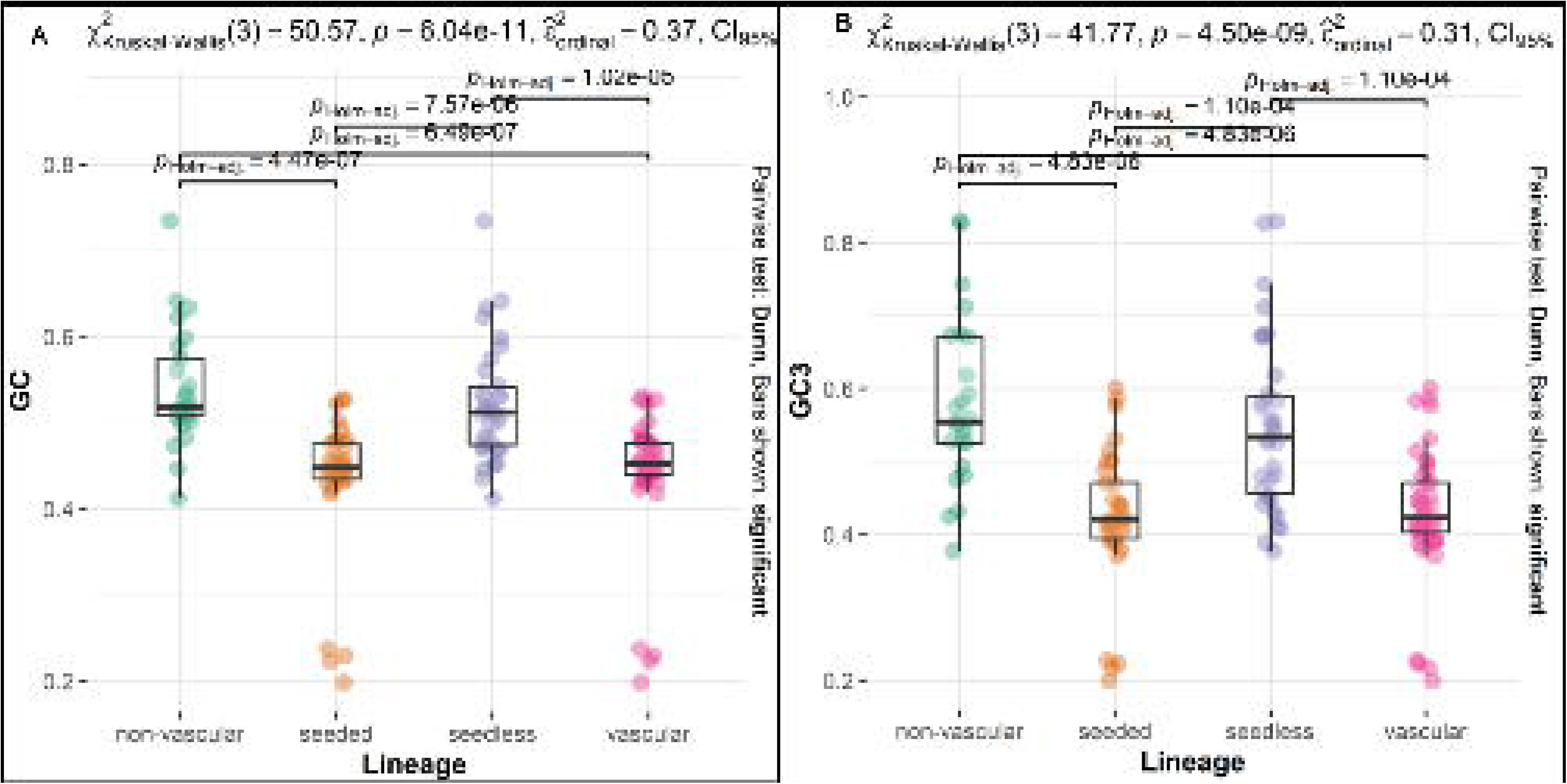
Variations in the pattern of GC-content (A) and GC3-content (B) with respect to vascularization and seed production.

### CUB is relatively stronger in algae than other lineages

The CUB relatively appears to be more strong in algae (mean ENC = 49.29) than those of bryophytes (mean ENC = 53.52), pteridophytes (mean ENC = 53.69), gymnosperms (mean ENC = 51.85), dicots (mean ENC = 52.27), and monocots (mean ENC = 51.53) (Figure 2A). *Chlamydomonas reinhardtii* was observed to exhibit the highest codon bias (ENC = 36.47), followed by *Pyropia sp.* (ENC = 38.39). Other chlorophyte algae such as *Fritschiella tuberosa*, *Stigeoclonium helveticum*, and *Chlorella vulgaris* and the rhodophyte alga like *Lympha mucosa* also showed relatively higher bias. ENC value ranges from 20 (absolute bias) to 61 (no bias). An ENC value of <36 represents strong bias (He *et al.,* 2016). Thus, our analysis revealed that the CUB is moderately strong in algae, but weak in other lineages. We observed a relatively more pronounced variation in the CUB with respect to vascularization (ENC = 50.98 and 52.30 in non-vascular and vascular lineages respectively, with a mean difference value of 1.310) than that of seed production (ENC = 51.70 and 51.93 in seedless and seeded lineages respectively, with mean difference value of 0.225), however statistical significance was observed only with respect to seed-production (Figure 2B).

**Figure 2:**
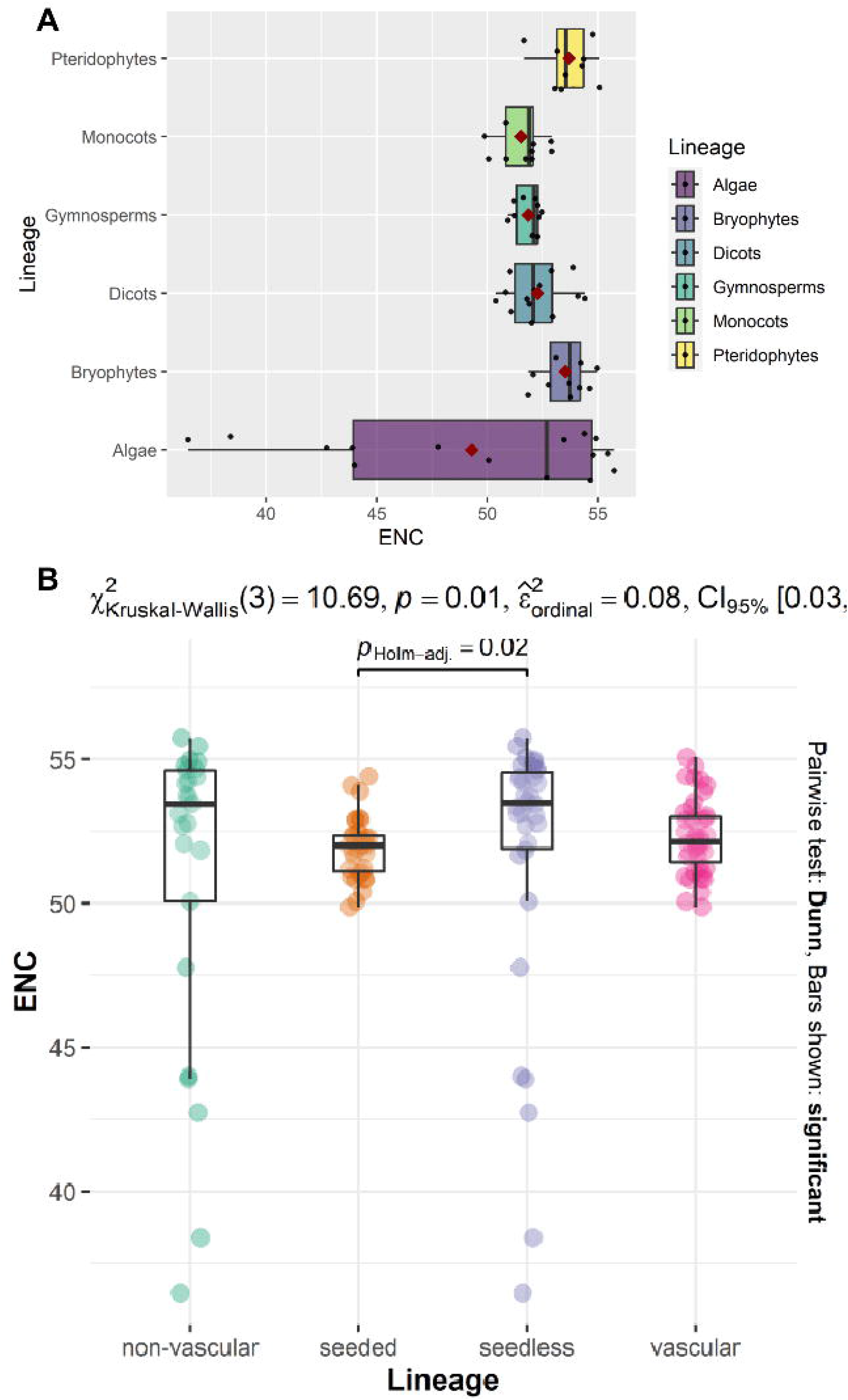
Summary of effective number of codon (ENC). (A) Representation of variation ENC values of different plant lineages. (B) ENC variation with respect to vascularization and seed production.

### Vascularization event might have influenced the codon usage patterns more prominently

In case of algae, all the optimal codons of the members of rhodophytes and the phaeophytes showed exclusive G/C-ending, whereas 82.92% of the optimal codons of chlorophyte members showed GC-ending (Supplementary file S1). The charophyte algae, however, showed relatively less proportion (53%) of GC-ending optimal codons. In the bryophytes, 80.2% of the optimal codons end with G/C, whereas the optimal codons were mostly AU-ending in the pteridophytes, gymnosperms, and dicots. However, all the optimal codons of *Selaginella rupestris* and *Wollemia nobilis*, and around 77% in *Ranunculus scleratus* exhibit GC-ending. In comparison to dicots, the monocots have a greater percentage of G/C-ending optimal codons. In contrast to other monocots, *Tradescantia pallida* exclusively shows AU-ending, whereas, in *Dioscorea prahensilis*, 93.33% of its optimal codons exhibit AU-ending. The RSCU is a measure of the CUB. RSCU value of 1 denotes no bias, whereas the values >1 indicate bias. Further, RSCU < 0.75 represent avoided cocoons (Guan *et al.,* 2018). Based on this criteria, the proportion of the avoided codons ending with AU was found to be relatively greater in algae and bryophytes. The pteridophytes, gymnosperms, dicots, and monocots, however, have more GC-ending avoided codons (Supplementary file S2). Thus, in general, the vascular lineage tend to prefer AU-ending codons and avoid GC-ending codons, whereas the non-vascular lineages tend to prefer GC-ending coodns and avoid AU-ending codons (Supplementary file S1 and S2).

We used correspondence analysis (CA) and model-based clustering analysis to observe the variation in the codon usage pattern across the different plant lineages. Based on the average RSCU values of 59 synonymous codons, CA was carried out to ascertain how the evolutionary events of vascularization and seed production are related to codon choice. In CA, the variation explained by the first two axes was found to be 81.22% and 8.39% respectively. A symmetric biplot based on axes 1 and axes 2 (Figure 3) shows the global variation in the data. The distance between the points gives a measure of their similarity/dissimilarity. Points with similar profiles lie close on the factor map. The red triangles correspond to the different plant lineages, whereas the blue points correspond to the codons. In CA, the vascular lineages cluster distinctly from non-vascular lineages. In model-based clustering analysis, a search for the best model showed that EVE (ellipsoidal shape of equal volume and orientation) with two components (clusters) is the best fit model for our data, based on the Baseyan Information Criterion (BIC) (Figure 4A). The classification of each codon to be in a specific cluster is presented in supplementary file S3. There are two types of codons, the one ending with G/C, and the other ending with A/U. The cluster plot (Figure 4B) showed the segregation of the codons into two clusters viz cluster 1 and cluster 2. The cluster assignment of each codon (Supplementary file S3), further revealed that the AU-ending codons are highly related to cluster 1, whereas the GC-ending codons best fit to cluster 2. From the optimal and the avoided codon analysis, it was observed that the higher plants generally prefer A/U-ending codons and avoid G/C-ending codons, whereas the lower plant showed a reverse trend. To further investigate this relation, the EVE model-based clustering plot segregated the lineages into two distinct clusters, and revealed a clear separation of the vascular and non-vascular lineages (Figure 4C). The algae and bryophytes formed one cluster, whereas the pteridophytes, gymnosperms, dicots, and monocots formed the other cluster. The cluster plot showed that the pteridophytes, gymnosperms, and dicots are closer to each other than the monocots. It is obvious because the monocots exhibit more GC-rich codons than dicots, so their proportion of the preferred G/C-ending codons is greater than those of the other vascular lineages, rendering them slightly distinct from other vascular lineages. Further, the density plot (Figure 5) generated through the model-based clustering, showed a close association of pteridophytes, gymnosperms, dicots, and monocots in codon choice. Similarly, the non-vascular lineages are more closely related to each other, than to the vascular lineages. Based on the results from the CA and model-based clustering, it, therefore, appears that the divergence of plants with respect to vascularization might have influenced the evolution of the codon usage pattern more prominently than the seed production. The statistical analysis through Kruskal-Wallis test, however, revealed no significant differences in codon usage pattern between the differently clustered lineages in CA and cluster analysis, when all the codons where considered together. Since, non-vascular lineages tend to prefer GC-ending codons in contrast to AU-ending codons by the vascular lineages (supplementary file S1), we therefore performed Kruskal-Wallis test based on the AU- and GC-ending codons, which showed significant differences in the mean RSCU values between the vascular and non-vascular lineages (Figure 6). There is a significant difference of 0.27 and 0.29 in the mean RSCU of AU-ending and GC-ending codons, respectively between the vascular and non-vascular lineages. Therefore, the clustering pattern obtained through CA and model based clustering, which revealed differences in codon usage patterns of vascular and non-vascular lineages can be attributed to their differential preference of AU- and GC-ending coodns.

**Figure 3:**
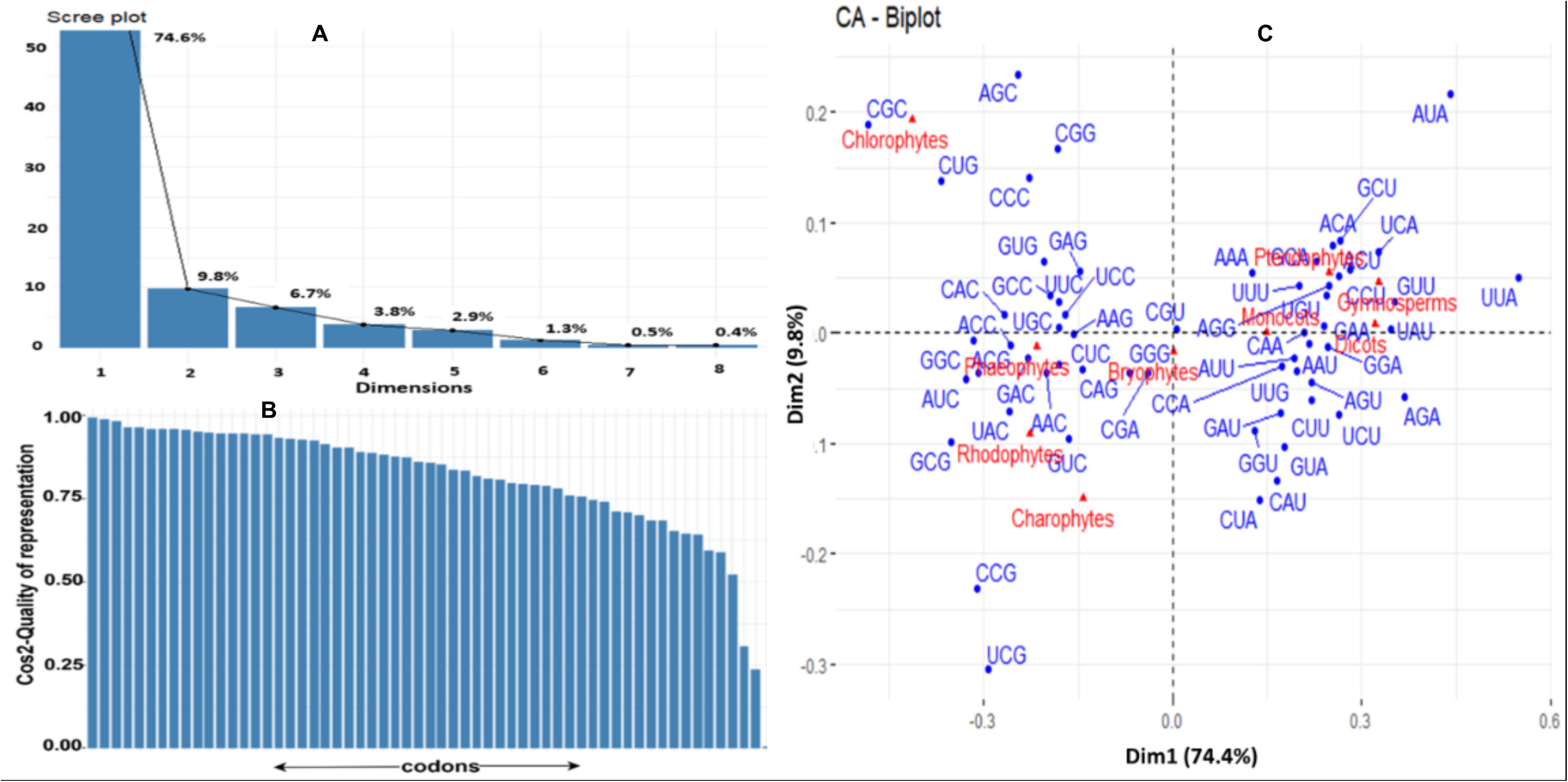
Correspondence analysis based on 59 synonymous codons. (A) Shows the scree plot representing relative contribution of different axis in explaining the variance. (B) Shows the cos2 (quality of representation) of variables on dimensions 1 and 2. (C) Represents symetric plot showing scatter of both codons and different plant lineages.

**Figure 4:**
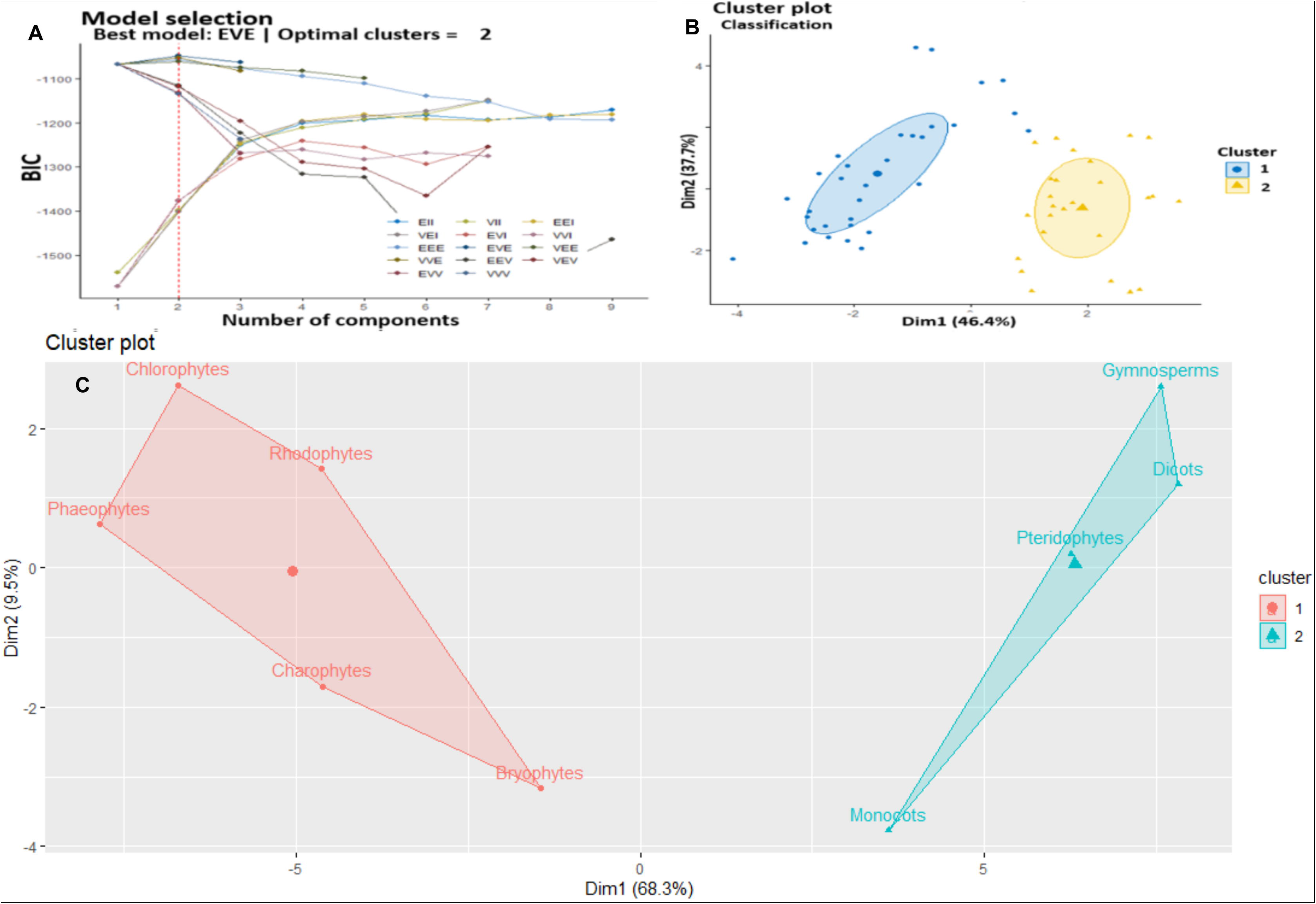
Details of model based clustering analysis. (A) Represents the results of the model selection. (B) Shows the classification of codons into their respective clusters. (C) Cluster plot showing the separation of different lineages into two clusters.

**Figure 5:**
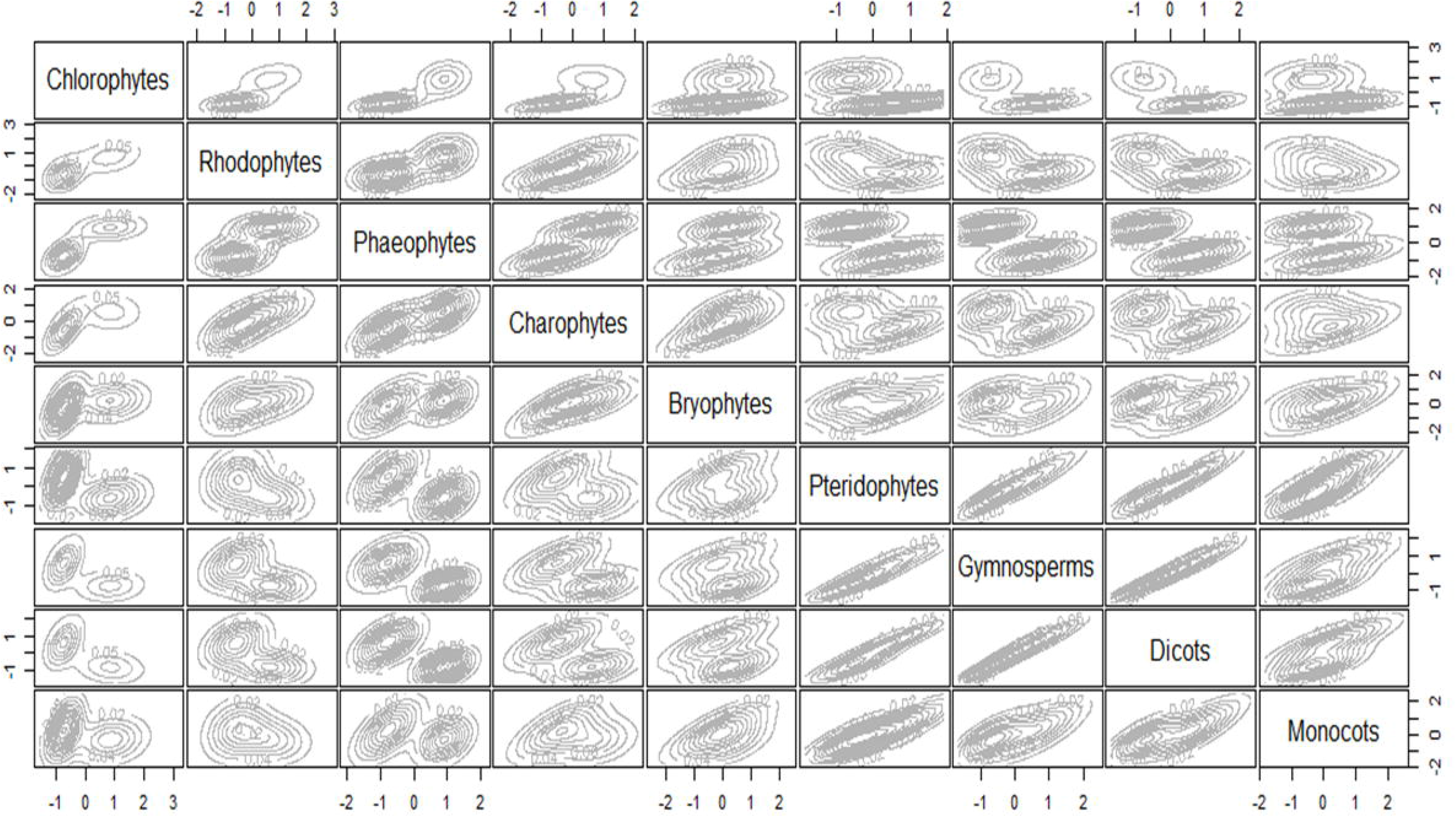
Density plot showing the relationship of different lineages with each other based on their mean RSCU values. The scale for each row and each column is based on the transformed RSCU values. More linear the density plot is, the more closer relationship between the groups.

**Figure 6:**
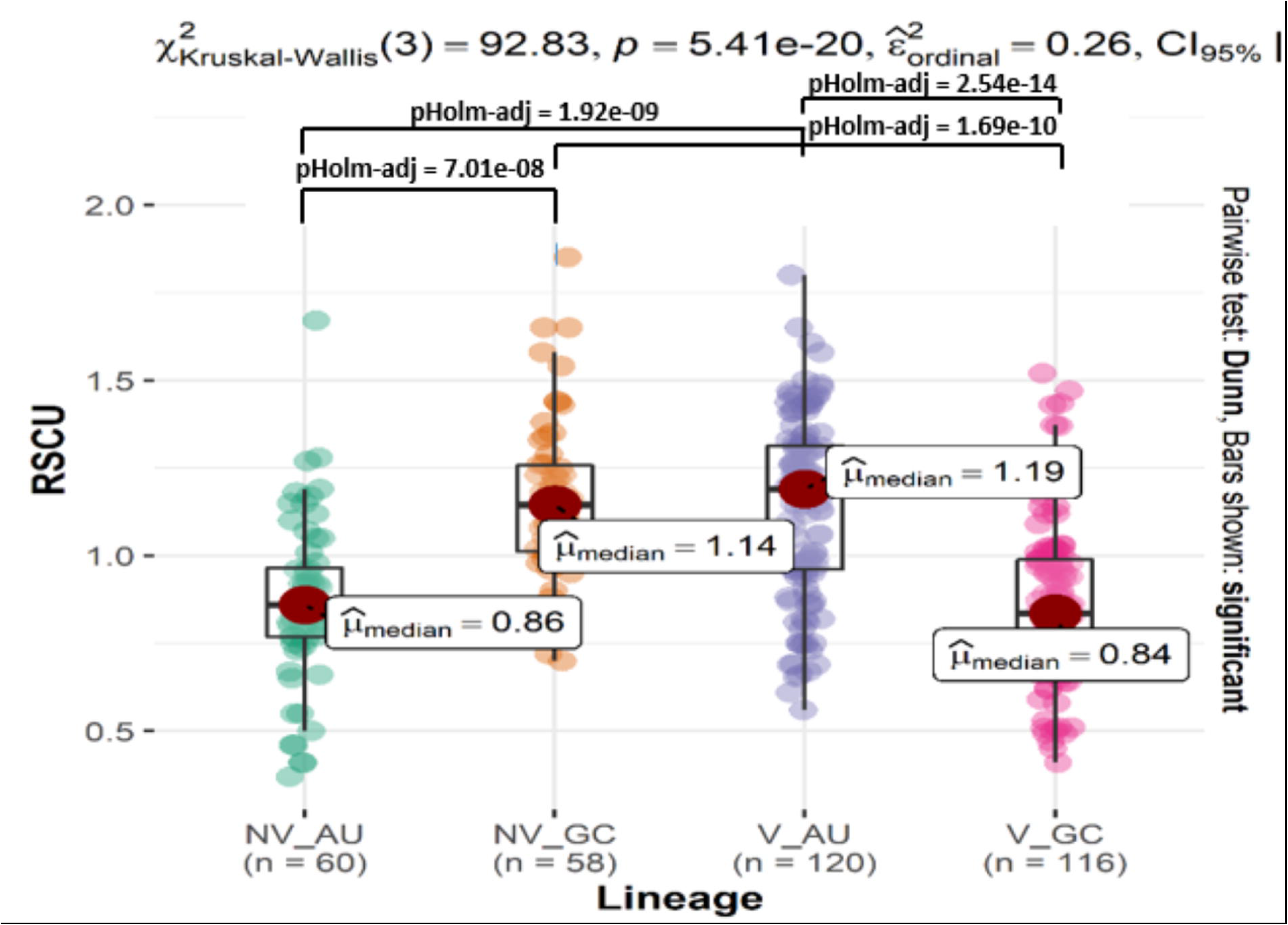
Representation of variation pattern in AU- and GC-ending codons between vascular and non-vascular lineages.

### Selection constraint predominates over mutational pressure in determining CUB

Estimation of relative influence of mutation and selection was carried out through neutrality plot analysis. The neutrality plots were generated by plotting the GC12 on the y-axis and the GC3 on the x-axis. GC3 mutational pressure drives the CUB if there is a significant correlation between the GC12 and the GC3, and the regression slope is close to one. As the slope approaches to horizontal or zero, selection becomes the dominant force in determining the CUB (Wu *et al.,* 2015). The slopes of the regression equation of chlorophytes, rhodophytes, phaeophytes, charophytes, bryophytes, pteridophytes, gymnosperms, dicots, and monocots were 0.154, 0.296, 0.177, 0.240, 0.124, 0.097, 0.088, 0.098, and 0.214 respectively (Supplementary files S4-S5). The relative neutrality of 0.154 in chlorophytes shows that the mutational pressure accounts up to 15.4% in determining the CUB, whereas the relative influence of selection is 84.6% (Chakraborty et al. 2017a, 2017b). Thus, based on the relative neutralities, it was observed that the selection plays a dominant role in all the lineages than that of the mutation in determining the CUB. In general, the non-vascular lineages have relatively higher mutational influence of 19.8% (average neutrality of 0.198) than those of vascular lineages 12.4% (average neutrality of 0.124).

### Dinucleotide bias and Codon pair bias (CPB) analysis

CUB can be influenced by dinucleotide bias (Chiusano et al. 2000; Karlin and Burge 1995). The frequencies of the individual 16 possible dinucleotides were calculated, followed by evaluation of their relative abundance (odds ratio) (Supplementary file S6). The dinucleotides CpG and TpA were under-represented in all the lineages, so can be considered as avoided dinucleotide markers. Under-representation of CpG was also observed in most virus hosts by Di Giallonardo *et al*., (2017). The dinucleotide TpG was found over-represeneted in all the lineages, except algae. We observed a clear distinction in the usage pattern of the dinucleotides ApA and CpA with respect to the vascularization and seed production (Figure 7). These dinucleotides were found over-represented in the non-vascular and seedless lineages and under-represented in the vascular and seed lineages. Further, the dinucleotides ApG and TpT were under-represented, whereas CpC and GpC were over-represented only in seed plants.

**Figure 7:**
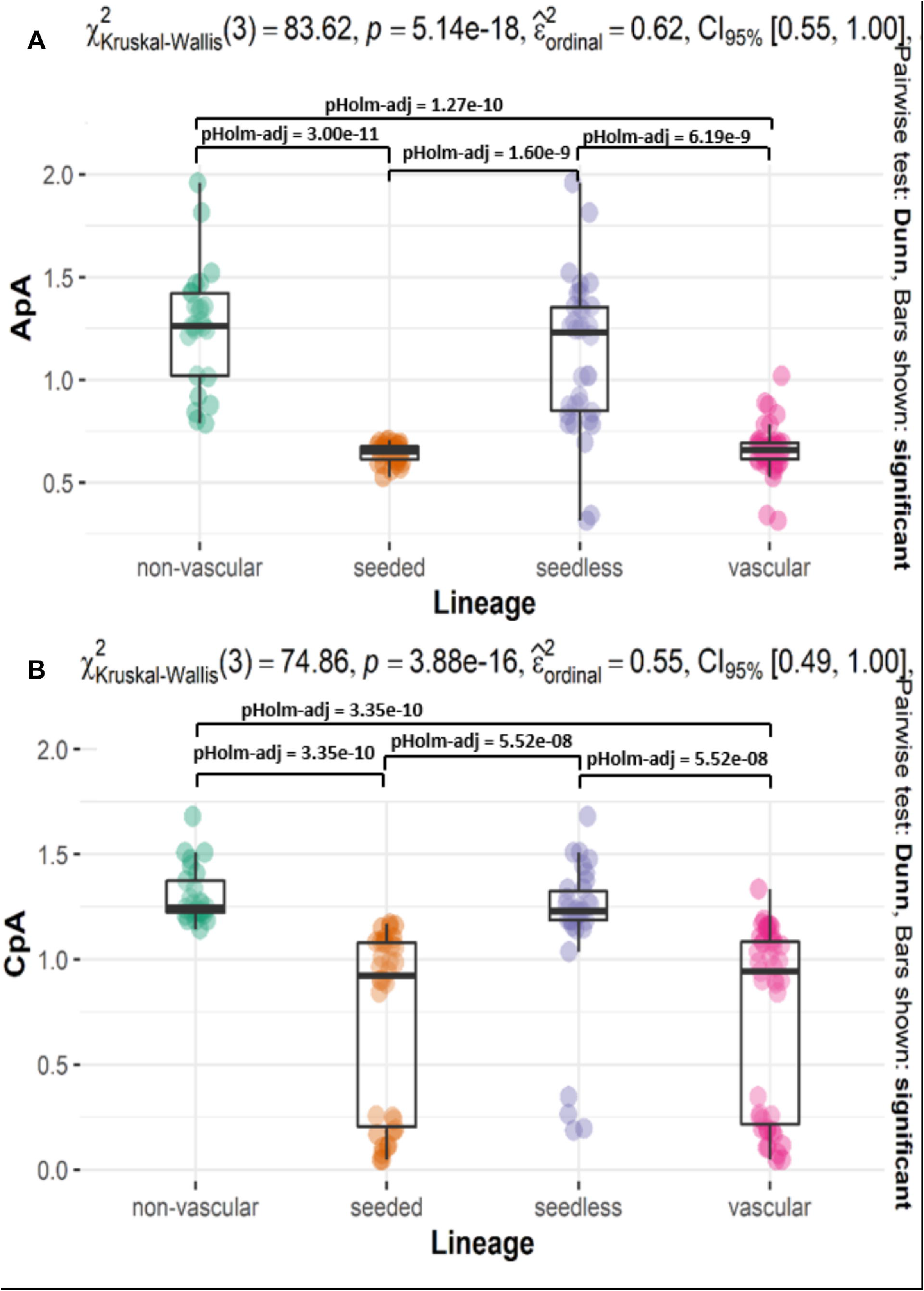
Representation of variation pattern in the dinucleotides ApA (A) and CpA (B) with respect to vascularization and seed production. Y-axis represents the odds ratio.

The members of a neighbouring codon pair may have a strong association and can be used more frequently than expected, thereby constituting a biased usage of the codon pair known as CPB. The later may affect the CUB. We observed the mean CPB values of -0.049, -0.052, -0.031, - 0.019, -0.032 and -0.041, -0.035, -0.038, and -0.052 for chlorophytes, charophytes, rhodophytes, phaeophytes, bryophytes, pteridophytes, gymnosperms, dicots, and monocots respectively. Based on the CPS values, the overrepresented codon pairs (CPS > 0) and the underrepresented codon pairs (CPS < 0) were identified, however, each lineage possessed more number of under-represented codon pairs than over-represenetd ones, due to which the mean CPB of each lineage was lower (Supplementary file S7). Since, the dinucleotide bias is linked with the CPB (Shen et al. 2015), the codon pairs were further analyzed to assess any dinucleotide bias. We observed that the dinucleotide CpA was over-represented in the over-represented codon pairs and under-represented in under-represented codon pairs. The dinucleotide CpG was over-represented only in the under-represented codon pairs. Further, GpA was under-represented both in the over-represented and the under-represented codon pairs. At the III-I positions of the juxtaposed codons, similar results were obtained for the dinucleotides CpA and GpA, whereas the CpG was under-represented and the ApC was over-represented in the over-represented codon pairs.

### Evolutionary related lineages have similar Codon pair Bias

A comparison of the CPSs across different lineages revealed that the vascular lineages have a strong relatioship with each other than with the non-vascular lineages. Similarly, the non-vascular lineages have a strong correlation with each other, but not as strong as seen among the vascular lineages (Figure 8). It is evident that the correlation of CPSs between the vascular lineages becomes weaker as the phylogenetic distance between them increases. Among the non-vascular lineages, the charophytes show a stronger relationship with the chlorophytes than those of the rhodophytes and the phaeophytes. Thus, in general, the phylogenetically closer lineages have a similar pattern in the choice of codon pairs. In our analysis, since the CPSs of phylogenetically related groups appear to be more similar, it generally reflects their common evolutionary history in addition to the common forces shaping their codon pairing.

**Figure 8:**
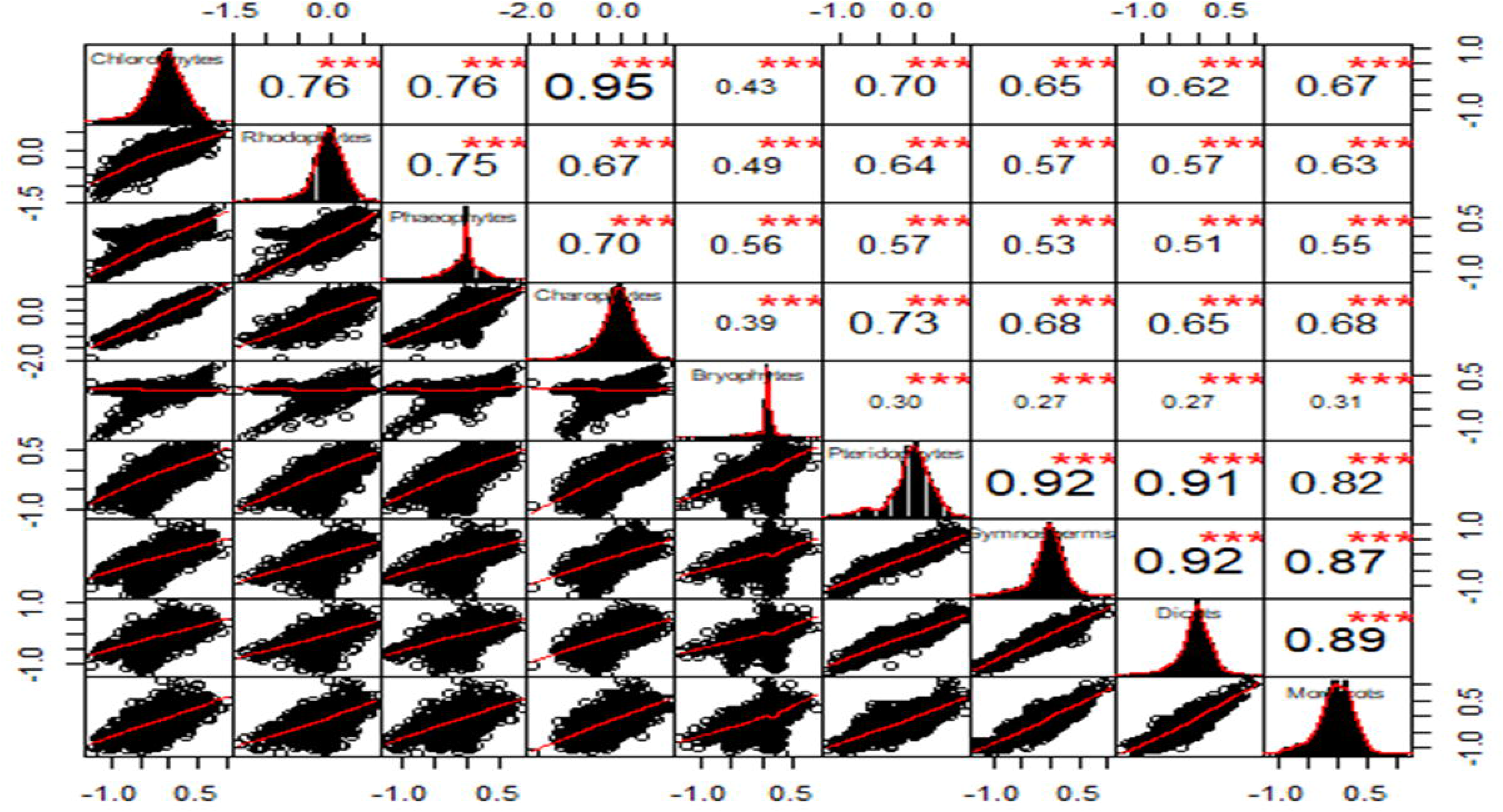
Correlation plot of CPS among different lineages. Histograms (black bars in the diagonal boxes), density plots (red curves in diagonal boxes), and regression plots (boxes in lower diagonal with black dots representing the scatter plot and red lines indicate regression slope). The numbers in the boxes (upper diagonal) represent correlation coefficients. The significance of the correlation is represented by red stars. X- and Y-axis of each box are based on CPSs values.

## Discussion

The variation of codon usage pattern in response to evolution of organisms is still an intriguing question to answer. The relative impact of the determining forces of CUB may vary in different plant lineages. CUB is not widely studied in plants and a comprehensive comparative analysis among the different plant lineages is lacking. The purpose of this study was to investigate the variations in the general pattern of codon usage and to evaluate the relative influence of mutation and selection in determining the CUB. Moreover, the relation of codon usage with respect to evolutionary events of vascularization and seed production was focused. We found that the CUB is moderate in algae and weaker in other plant lineages. The ENC value obtained in this study for bryophytes is comparable to the model moss, *Physcomitrella patens* (Szövényi et al. 2017). Further, in *G. biloba* and *T. contorta*, He *et al.,* (2016) and Majeed *et al.,* (2020) found ENC values similar to our study for the gymnosperms respectively. The ENC values of the vast majority of the unigenes were found to be high indicating very weak bias in the codon choice. This suggests that during protein synthesis, almost all types of amino acids are used by these lineages, except chlorophytes and rhodophytes, and a preference for the specific codons has not evolved strongly in their genomes. However, in chlorophytes and rhodophytes, the CUB may strongly govern the selection of specific codons (Guan *et al.,* 2018). Based on the mitochondrial genomes, a similar comparatively higher bias was seen for algae in comparison to bryophytes, monocots and dicots, which show relatively similar ENC values (Wang et al. 2011). It is believed that the CUB regulates the rate of protein synthesis. Frequent codons typically accumulate in highly expressed genes and accelerate translation rate because of the abundance of their corresponding tRNAs, whereas rare codons accumulate in low expressed genes and slow down the translation rate. Besides, CUB regulates translational accuracy, transcription, and co-translational folding (Xu et al. 2021). In addition, CUB influences mRNA levels through transcription in a translation-independent manner through multiple transcriptional regulatory mechanisms (Zhau et al. 2021). Our study did not focus on evaluating the relationship between the CUB and gene expression regulation. It would be interesting to explore this relationship in diverse plant lineages.

We found the highest mean GC-content in the codons of chlorophytes (58.84%), followed by rhodophytes (58.49%), and phaeophytes (56.72%), and the lowest in dicots (39%). A high GC-content (63.6%) of the genome of marine red algae, *Pyropia yezoensis* was observed (Nakamura et al. 2013). Further, the GC-content of the EST-sequences of this species was estimated to be 65.2%. Interestingly, the codons with high GC-content at the third position were frequently used. Similar to our results, the GC-content of *P. yezoensis* at the third codon position was highest (79.4%) followed by GC1 (62.2%) and GC2 (45%) (Nikaido et al. 2000). This suggests that like the genome of *P. yezoensis*, the algal genomes, in general, have been exposed to high GC-pressure during the evolution. Based on the transcriptomic sequences, the GC-content of the Liverwort, *Marchantia polymorpha* was estimated to be 48.6%, which is similar to the mean GC-content (47.07%) of the pteridophytes found in our study, but lower than the lycophyte, *Selaginella* (52.5%) and green algae *Chlamydomonas* (64.5%) (Sharma et al. 2014). The mean average GC-content of the pteridophytes in our study is higher than that of the model moss, *Physcomitrella patens* (Szövényi et al. 2017), but similar to the average detected GC-content in overall land plants (Singh et al. 2016). The average GC-content of *G. biloba* (44.88%) and it’s higher GC1-content (51.39%) than GC2 (40.88%) and GC3 (39.02%) found by He *et al.,* (2016) is in agreement with our finding for gymnosperms (GC=44%, GC1=51.20%, GC2= 40.08% and GC3=41.3%). A higher GC-content in monocots than dicots is due to the presence of more GC-rich gene class and having no or few short introns in the former (Carels and Bernardi 2000). These differences might govern the corresponding differences in codon usage found in this study, which are in agreement with previous studies (Fennoy and Bailey-Serres 1993; Sundararajan et al. 2016). In general, the GC-content of higher plants, (vascular seedless and vascular seeded) is higher than those of the lower plants like bryophytes and heterogeneous algal groups (Figure 1). It, therefore, appears that the evolutionary events of vascularization and seed production have selected the plant species for relatively lower GC-content.

Although there were slight variations among the individual species, the optimal codons of the vascular plant lineages, pteridophytes, gymnosperms, monocots, and dicots show preference for AU-ending optimal codons. Monocots however have more GC-ending optimal codons than dicots. On the other side, all the non-vascular lineages showed a preference for GC-ending optimal codons. Our analysis showed a statistically significant difference in the optimal codon usage between the vascular and non-vascular, as well as between the seedless and the seed plants. However, the difference was higher with respect to the vascularization than that of the seed production. This suggested that although both the events generated significant variations in the optimal codon choice, the relative influence of the vascularization is sharper than that of the seed production. It, therefore, appears that the pattern of optimal codon usage has evolved more strongly with vascularization in the evolutionary history of plants. In contrast to the optimal codons, the pattern of the avoided codon usage followed a reverse trend, with higher lineages generally showing GC-ending and lower lineages generally showing AU-ending. Like the optimal codons, the vascularization event turned to be more significant than that of the seed production, in governing the pattern of avoided codons. Our findings are corroborated by the observation of optimal algal codons biased towards GC-ending, with codons of *Griffithsia okiensis* and *Chondrus crispus* having mostly C-ending, whereas those of *Porphyra yezoensis* having mostly G-ending (Lee et al. 2007). Based on the analysis of CUB in mitochondrial genes, plants, in general, were found to have frequent AU-ending preferred codons, with the strongest preference in bryophytes and weakest in land plants (Xu et al. 2015). However, this preference was strongly linked to intron number in nuclear genes (Qin et al. 2013). The A/U-ending preferred codons were also found more than G/C-ending in the dicot species like Chinese berry (Feng et al. 2013) and 19 citrus species including *Citrus sinensis*, *C. clementina*, *C. reticulata*, and *Poncirus trifoliata* (Ahmad et al. 2013). Further, the gymnosperms, *G. biloba* (He *et al.,* 2016) and *T. contorta* (*Majeed et al., 2020*) were also found with a higher proportion of A/U ending preferred and G/C ending avoided codons. We observed that monocots show the predominance of G/C-ending codons. *Oryza sativa*, *Zea mays*, *Triticum aestivum*, *Hordeum vulgare* were shown to have GC-ending preference, thereby corroborating our results (Kawabe and Miyashita 2003; Liu and Xue 2005b). This distinction of monocots from dicots is also observed by Campbell and Gowri (1990). The preferential G/C-ending codons of monocots can be attributed to the presence of two types of genes viz the normal genes like those in dicots and the genes having high G/C bias (Brinkmann et al. 1987; Campbell and Gowri 1990; Jansson et al. 1994). It appears that those G/C biased genes predominate in monocots. In this study, we found that gymnosperms exhibit the highest preference for A/U-ending codons, which is in agreement with He *et al.,* (2016). The overall GC-bias of a genome or genomic region is highly correlated with GC3. G/C-ending codons should increase in frequency with increasing GC-bias, whereas that of A/U-ending codons should decrease (Kliman and Bernal 2005). Thus, the preference for GC-ending codons in algae groups and bryophytes can be linked to its high GC-content.

Our study showed that natural selection predominates over mutational pressure in determining CUB in all the lineages. However, the relative impact of the mutation decreases, whereas that of the selection increases from algae to angiosperms. Shields et al. (1988) first showed the involvement of natural selection in *Drosophila melanogaster,* which was eventually also revealed in many other studies (Akashi 1995; Akashi and Schaeffer 1997; McVean and Vieira 2001). Further, the selection strength was found to be weaker in charophyte algae (Wang et al. 2011). Moreover, the studies utilizing the coding sequences found stronger selection constraints than mutational pressure in determining CUB in higher plants like *G. biloba* (He *et al.,* 2016) and *T. contorta* (Majeed et al. 2020). In agreement to our study, natural selection played a major role in determining CUB in four pteridophytes, *Calypogeia arguta*, *C. integristipula*, *C. neogaea*, and *C. suecica* (Das et al. 2019), *Bombyx mori* (Jia et al. 2015), *Oryza sativa*, *Zea mays*, *Triticum aestivum* and *Arabidopsis thaliana* (Liu and Xue, 2005), *Triticum aestivum*, *Oryza sativa*, *Zea mays*, *Hordeum vulgare*, *Arabidopsis thaliana*, *Nicotiana tobaccum*, and *Pisum sativum* (Kawabe and Miyashita 2003). In addition to mutation and selection, genetic drift and effective population size are also thought to affect CUB. Species having low effective population sizes exhibit less variability and reduced effectiveness of selection (Charlesworth 2009). In organisms with insufficient effective population size and long generation times, selection is relatively weaker (Rocha 2004; Lynch et al. 2006; Subramanian 2008; Wang et al. 2011). Even though the CUB and selection were significantly correlated, effective population size did not predict variations in CUB in mammals, suggesting that the distribution of selection coefficients varies across species independently of effective population size, confounding any straightforward link to CUB (Kessler and Dean 2014). The authors argued that CUB is influenced by diverse factors and species divergence in any of these factors, plus the lack of appropriate data for parsing out different forces affecting CUB obscure in establishing any direct link between CUB and effective population size. Further, species with small population sizes have increased recombination rates. Possibly, the reduced effective population size may be counterbalanced by enhanced selective efficiency gained by elevated rates of recombination, thereby making straightforward predictions about the relation between population size and CUB more complicated (Kessler and Dean 2014). Due to unavailability of the required data to directly establish a link between CUB with effective population size and drift, we restricted our analysis to mutation anmd selection.

Dinucleotide bias may affect the pattern of codon usage (Karlin and Burge, 1995; Chiusano *et al*., 2000). The relative abundance of 16 dinucleotides in the coding sequences was evaluated to assess the dinucleotides affecting the codon usage patterns. Our observation of over-representation of TpG (except algae) and under-representation of CpG in all plant lineages is corroborated by (Chakraborty et al. 2017b). CpG dinucleotides are significantly suppressed in the genomes of most RNA and DNA viruses of vertebrate species (Karlin et al. 1994). They were also found the most under-represented dinucleotides in vertebrate genomes (Kunec and Osterrieder 2016). Similar to our results, they also found TpG most over-represented in vertebrates. Our study revealed that the dinucleotides ApA and CpA were responsive to vascularization. Another aspect of the codon bias involves the relation between the neighbouring codons. Like the non-random usage of the synonymous codons, similarly and independently of this bias, the juxtaposed codons are also not randomly used. Some codon pairs are used either more or less frequently than expected from their individual frequencies (Coleman et al. 2008; Kunec and Osterrieder 2016). This is known as codon pair bias (CPB) and is a stable character of a species. CPB is evaluated by estimating CPS. A negative value of CPS indicates that the codon pair is underrepresented, whereas a positive value reveals that the codon pair is overrepresented(Kunec and Osterrieder 2016). Our analysis showed lower CPB values in all the lineages, which can be attributed to the higher proportion of the underrepresented codon pairs than the overrepresented codon pairs (Kunec and Osterrieder 2016). CPB together with CUB represents two layers of codon bias. It must be noted that while the selection on CPB is context-dependent, the selection on CUB is context-independent. The codon context is the relationship between two adjacent codons (Chu and Wei 2021). The codon pair bias may be influenced by the dinucleotide bias (Shen et al. 2015). We observed that the dinucleotide CpA was over-represented in the over-represented codon pairs and under-represented in under-represented codon pairs, even at the codon junctions. This similar representation in the usage of the CpA and their parent codon pairs indicate that CpA may significantly influence the codon pair choice. Our analysis showed that only a small proportion of the codon pairs were responsive to the evolutionary events of either the vascularization or the seed production. The dinucleotide analysis of such codon pairs revealed ApA and CpC were responsive to the evolutionary event of the seed production, whereas GpC and TpG were responsive to the evolutionary event of vascularization. In this study we estimated CPB of different plant lineages and found that phylogenetically distinct lineages show different codon pair preferences and more related lineages exhibit similar CPB, reflecting their common evolutionary history and forces shaping codon pairing in their genomes. Similar results were also obtained by Kunec and Osterrieder, (2016) among the different species of vertebrates and arthropods. Our analysis was based on the reconstructed transcripts, as such it was not possible to appreciate the difference between non-transcribed sequences or introns and exons, whcich mighnt affect the dinucleotide bias (Porceddu and Camiolo 2011; Camiolo et al. 2015).

This study utilized RNAseq data to evaluate the pattern of CUB across different plant lineages, determining forces for CUB and contrasted the CUP pattern between seed vs seedless and vascular vs non-vascular plant lineages. The RNAseq data can further be utilized for SNP calling to perform population genetic analysis for CUB across these (Chu and Wei 2020) and RNA modification patterns (Chu and Wei 2021). Further, the variation in CUB patterns in different environments could be an interesting step to understand how different plant lineages manipulate their codon usage when subjected to a specific environment.

## Supplementary information

**S1:** Proportion of optimal codons in different plant lineages.

**S2:** Proportion of avoided codons in different plant lineages.

**S3:** Classification of codons into different clusters.

**S4:** Neutrality plots of chlorophytes (A), rhodophytes (B), phaeophytes (C), and charophytes (D).

**S5:** Neutrality plots of bryophytes (A), pteridophytes (B), gymnosperms (C), dicots (D), and monocots (E).

**S6:** Representation of bias in 16 dinucleotides across different plant lineages. The y-axis shows the odds ratio.

**S7:** Depiction CPB values of species of different plant lineages.

## Declarations

### Availability of data and material

N/A

### Code availability

N/A

### Authors’ contributions

PB concived and organized the over all study. AM performed the analysis and wrote the amnuscript. WR, AK,SD, JJ and AS peformed the computational studies. PB further edited and finalized the manuscript. All the authors have read and finalized the manuscript. The authors declare no conflict of interest.

### Additional declarations for articles in life science journals that report the results of studies involving humans and/or animals

N/A

### Ethics approval

N/A

## Acknowledgement

This study was supported by Central University of Punjab. Aasim Majeed acknowledges CSIR New Delhi for financial assistance during the PhD programme.

