## Supplementary figures and images for "Comprehensive Codon Usage Analysis Across Diverse Plant Lineages"

### Supplementary file S1

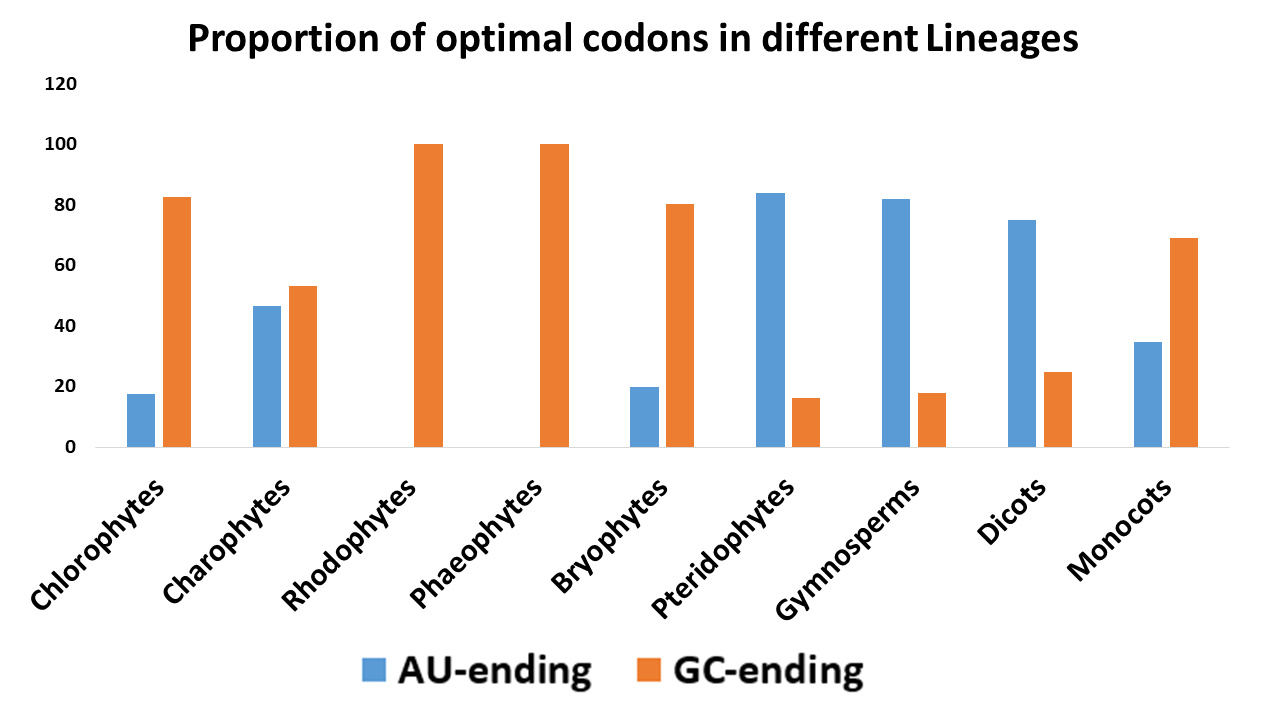

### Supplementary file S2

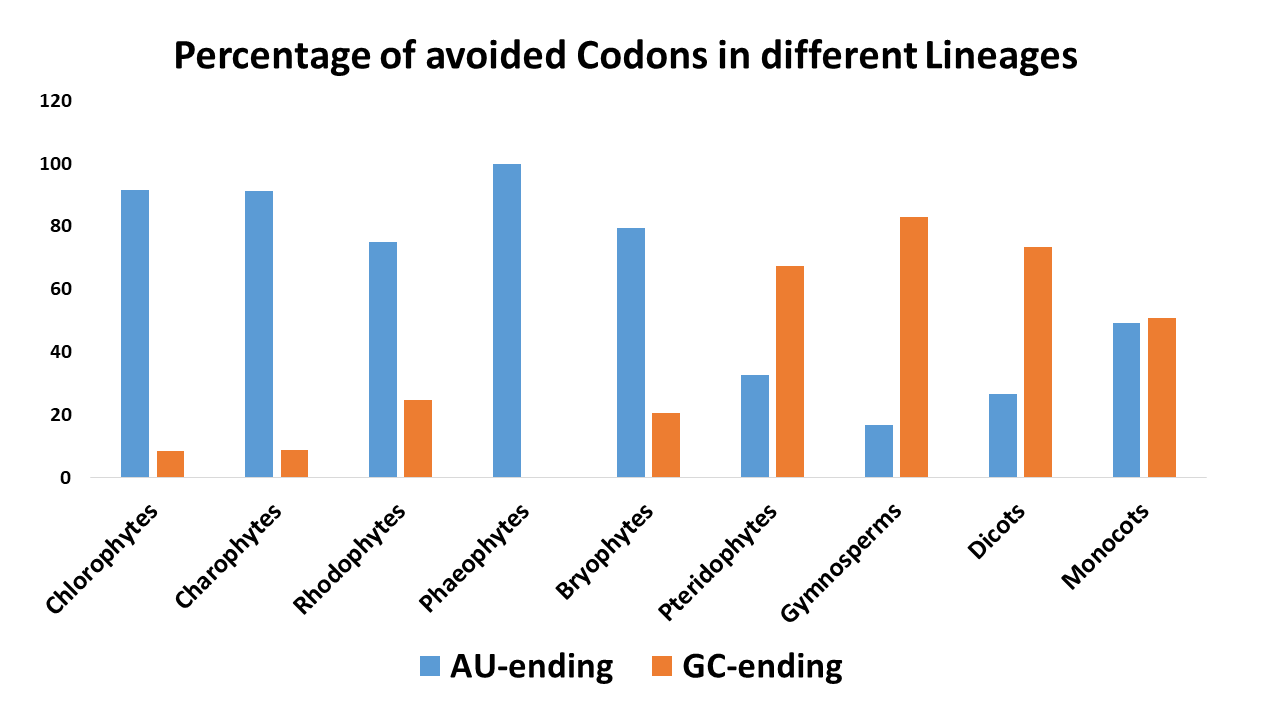

### Supplementary file S4

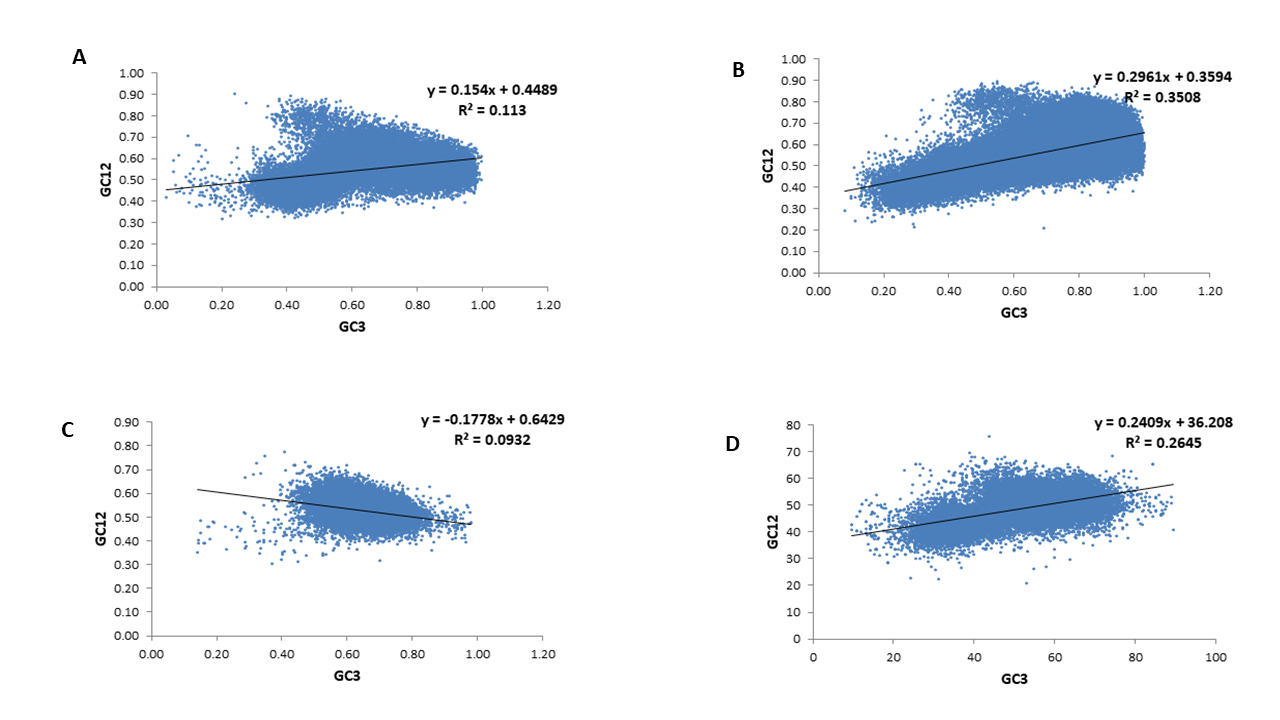

### Supplementary file S5

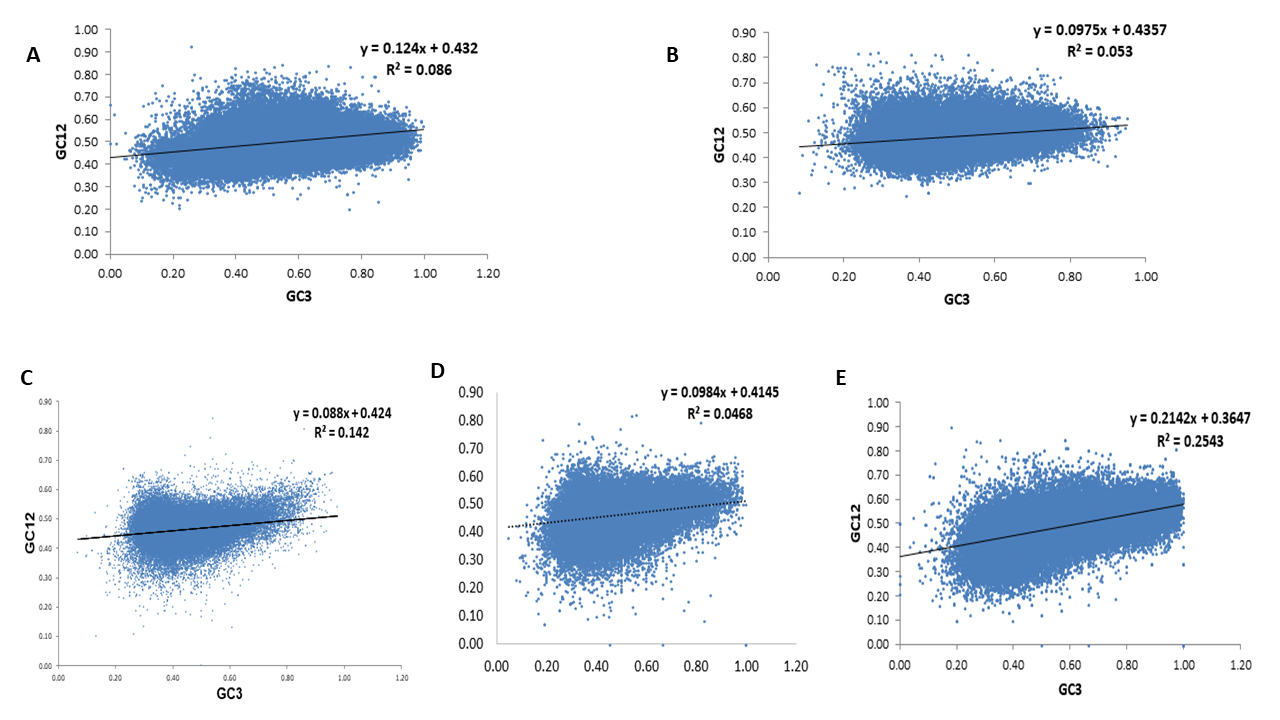

### Supplementary file S6

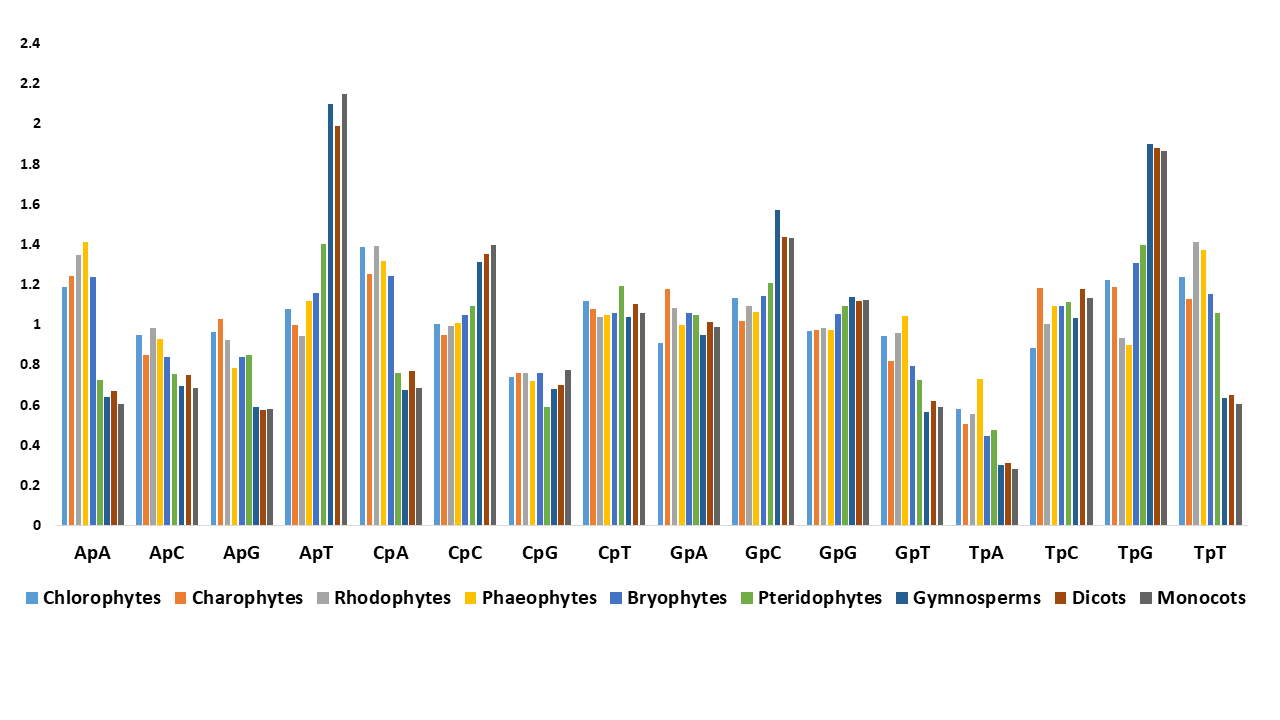

### Supplementary file S7

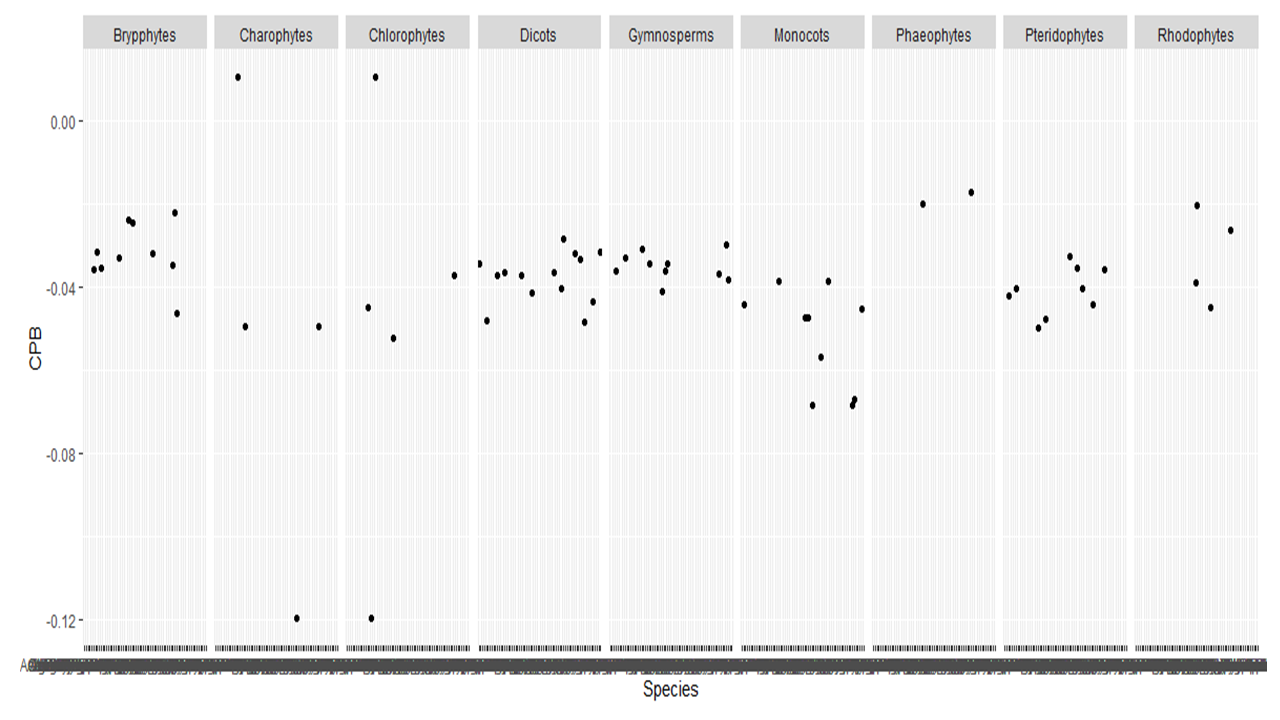
